# Synergistic information in the frontal cortex-striatal pathway

**DOI:** 10.1101/2021.06.18.449072

**Authors:** Ibrahim Alsolami, Takashi Handa, Tomoki Fukai

## Abstract

Across the cortico-basal ganglia circuit, the medial frontal cortex (MFC) communicates with the dorsal striatum (DS) during learning and planning. How these two brain regions communicate with each other is, however, not fully understood. Here we report the presence of synergistic information during information transfer across the frontal cortex-striatal pathway. Synergistic information emerges from the positive interaction of DS and MFC neurons and provides the DS with additional cortical information. This information is held latent in neuronal signals. To reveal it, we simultaneously record neuronal activities from the MFC and DS of rats trained on an outcome-based decision-making task and determined whether past neuronal activities of the DS positively influence communication rates. We detect a neuronal synergy that enables the MFC to boost its communication rate to the DS. Our results suggest that past neuronal activities of the DS help decode MFC signals. This ability is not attributed to the inherent autocorrelation of DS spiking activities.

## INTRODUCTION

The medial frontal cortex (MFC) and dorsal striatum (DS) work in tandem to bring about positive outcomes. Across the cortico-basal ganglia circuit, the MFC projects monosynaptically to the DS (Wilson, 1986; McGeorge and Faull, 1989), providing it with vital state information, such as a reward has been acquired or at times a lack thereof. In this neuronal circuit, deliberation and choice selection takes place during goal-directed behaviors (Lo and Wang, 2006). The MFC communicates error signals to the DS and is likely to adjust action by monitoring outcomes of prior decisions (Narayanan et al., 2013; Bonini et al., 2014; Hyman et al., 2017). Based on the information the DS receives from the MFC, it associates a specific selected action with a resultant outcome (Reynolds et al., 2001; Lauwereyns et al., 2002; Balleine et al., 2007; Nicola, 2007; Isomura et al., 2013; Nonomura et al., 2018). This is especially evident in laboratory studies, which demonstrate that DS-lesions in rats notably compromise their performance in action selection tasks (Skelin et al., 2014).

The activity of *in vivo* cortical neurons is highly irregular, and neuronal noise limits the amount of information that can be conveyed from the MFC to the DS. Yet the DS can still make adequate behavioral decisions with such irregular and noisy cortical inputs. This unique ability of the DS likely points to a decoding mechanism operating. In this study, we explore whether such a mechanism operates in the DS to decode MFC signals. Precisely, we determine whether past neuronal activities of the DS enable it to extract more information from MFC neuronal signals than otherwise without them.

Fundamentally, the more information the DS has from the MFC, the higher the chance an adequate decision is made. A key question therefore is: how much information is the DS extracting from MFC signals? To answer this, we simultaneously recorded neuronal signals from both the MFC and DS of rats voluntarily performing a left/right licking choice task. In this task, rats had to consistently switch spouts after a reward was withheld (Handa et al., 2021).

A central finding in this study is that past neuronal activities of the DS, in the millisecond regime, positively influence communication rates between the MFC and DS. This finding is especially intriguing because it hints at the existence of a consecutive decoding scheme at the DS, which uses past DS signals to better decode MFC signals. Crucially, the observation that past neuronal activities of the striatum contain information needed for subsequent actions is expected (Zhou et al., 2020; Handa et al., 2021). What is unexpected here, however, is that this information is synergistic—more than merely additive (equation 5). This finding suggests that the DS is not merely decoding instantaneous MFC signals but can also anticipate the state of the MFC.

## RESULTS

Twelve head-restrained rats were trained to lick either a left or right spout to acquire a reward allocated to one of the spouts (Figure 1A). The location of the rewarding spout was fixed during a trial block and was reversed after a rat accumulated a total of 10 rewards (Figure 1B). During this behavioral task, we simultaneously recorded neuronal activity from both the MFC and DS on the left hemisphere of rats with a pair of silicon probes. A retrograde tracer confirmed the projection from the recording site at the MFC to its corresponding recording site at the DS (Figure 2). In-depth details of our experiment are available in our recent report (Handa et al., 2021).

**Figure 1.**
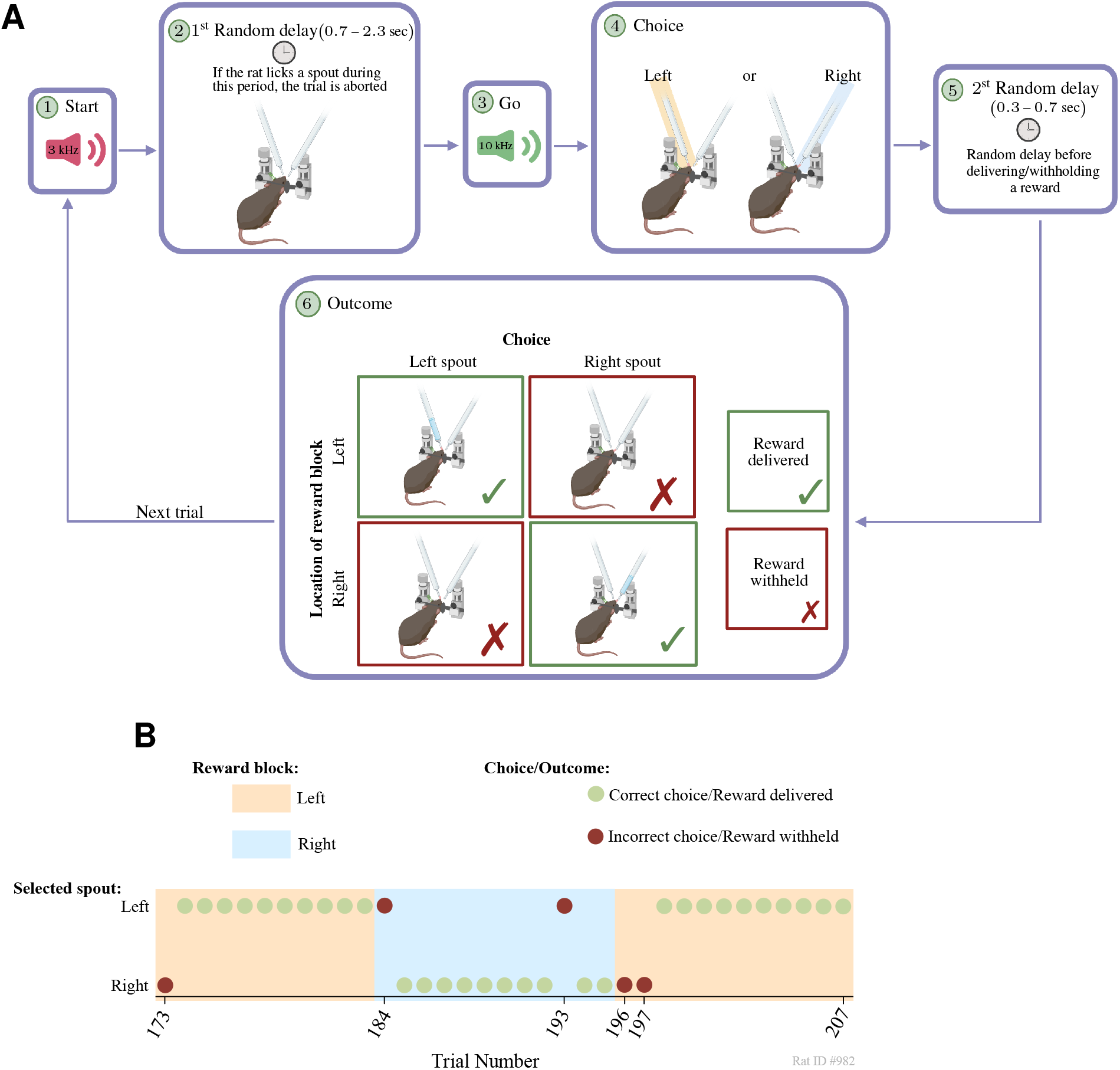
Experiment. (A) Schematic illustration of a single test trial: (1) A trial starts with a 3 kHz (1 sec) tone. (2) A random delay (0.7–2.3 sec) is then introduced. If a rat licks a spout during this delay period, the trial is aborted. (3) Next, a 10 kHz (0.1 sec) “Go” signal is presented. (4) After this signal, a rat selects either a left or right spout. (5) This is followed by a second random delay (0.3–0.7 sec). (6) Lastly, a reward is delivered if the selected spout location matches the reward block location; otherwise, a reward is withheld. After this step, a new trial begins. (B) An example from our behavioral experiment. First, the rat selects a spout (left/right). Then, a reward is delivered if the rat selects a spout location that matches the location (left/right) of a reward block. Otherwise, the reward is withheld. The location of the reward block is reversed after the rat accumulates a total of 10 rewards. Throughout the experiment, we provided no sensory feedback to rats regarding the location of rewards. In the above example, all the decisions made by the rat are correct except for trials #173, 184, 193, 196, and 197 (because the location of the selected spout does not match the location of the reward block). The incorrect choices made in trials #173, 184, and 196 are highly expected as rats tend to select the same rewarding spouts until it does not provide a reward. After which, rats tend to switch to an opposite spout, such as the decisions illustrated in trails #174, 185, 194, and 198 (that is to say, rats exhibit a win-stay/lose-shift strategy). In trial #197, however, the rat made a consecutive incorrect choice despite not gaining a reward in the previous trial #196; this case was not uncommon in our experiment. Additionally, in trial #193, the rat switched to an incorrect spout despite gaining a reward in the previous trial #192; while this case is uncommon, we observed it in our experiment.

**Figure 2.**
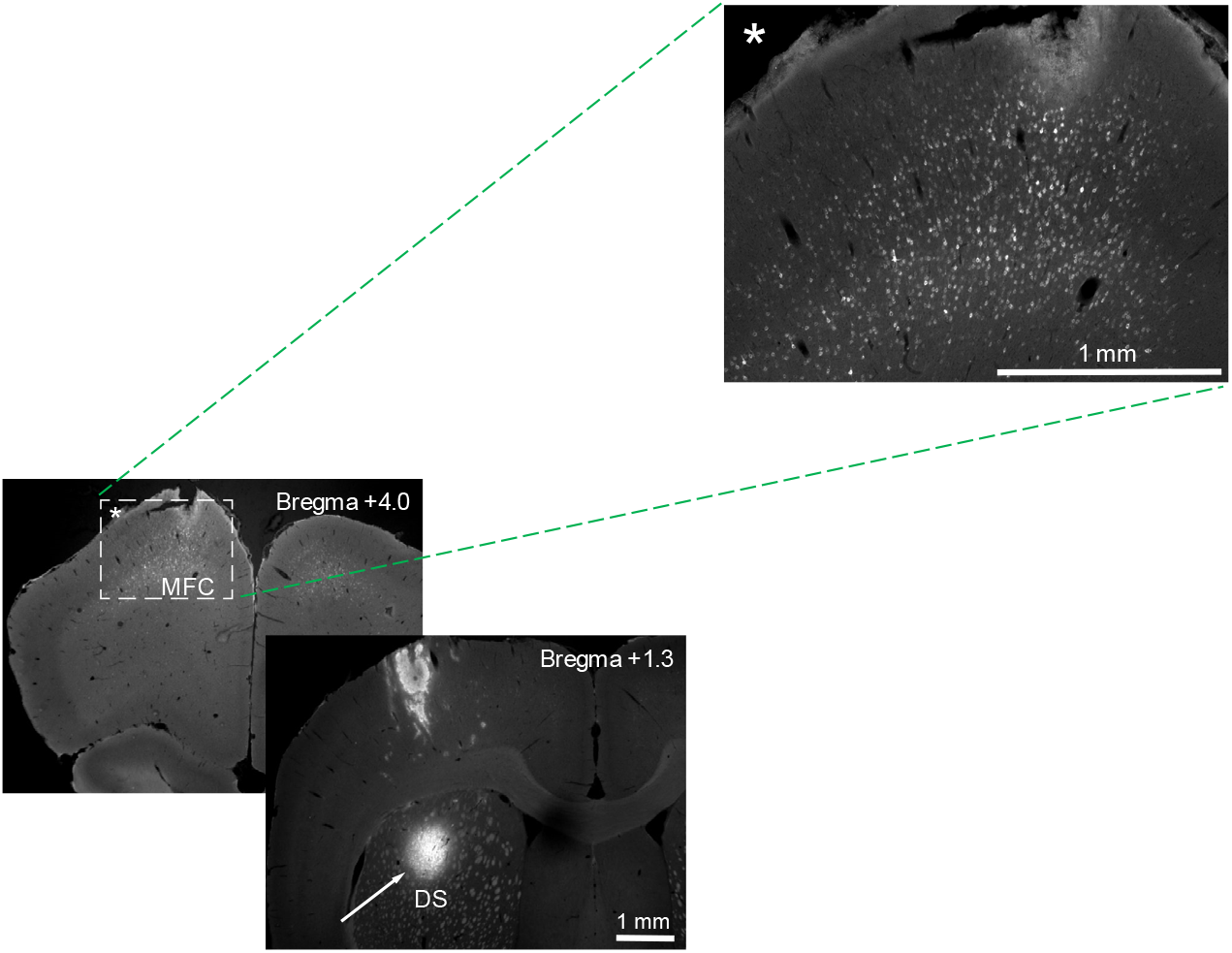
Fluorescent images of brain sections. Bottom left: Fluoro-gold labelled MFC neurons and injection site of tracer (Fluoro-Gold) at the DS. Top right: Expanded view of labelled MFC neurons in deep layers.

The remainder of this section is organized as follows: we first briefly review the theoretical background helpful for interpreting the neuronal data. We then present our experimental results, and discuss a frequent misconception regarding the seeming relationship between synergistic information and autocorrelation of DS neuronal activities.

### Synergistic Information and its Implications in Neural Communication

We quantify the amount of synergy forming between the MFC and DS from an information-theoretic standpoint. We briefly explain the information-theoretic measures employed in this study. In Figure 3, *X* is a random variable (RV) that counts the number of spikes generated by all MFC neurons within a time-window of duration *τ*_1_. Likewise, *Y* and *Z* are RVs that count the number of spikes generated by all DS neurons within a non-overlapping sliding time-window of duration *τ*_1_ and *τ*_2_, respectively. Herein, we evaluate the amount of synergistic information at the neural population level but not at that of single neurons, where large noise may arise due to sparse neuronal firing.

**Figure 3.**
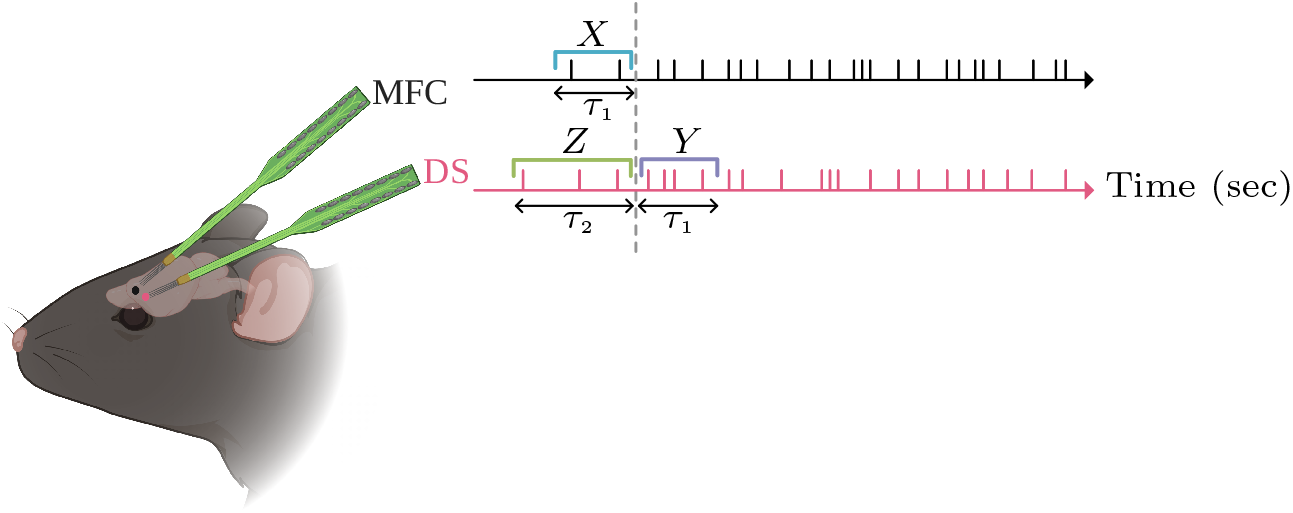
An illustration of the electrophysiological recording and random variables (RVs) involved. The MFC spike train displays the collective firing of all MFC neurons, rather than the firing of an individual MFC neuron, and likewise for the DS spike train. RV *X* counts the number of spikes generated by all MFC neurons within a time-window of duration *τ*_1_. Similarly, RVs *Y* and *Z* count the number of spikes generated by all DS neurons within a time-window of duration *τ*_1_ and *τ*_2_, respectively.

The mutual information quantifies the amount of information shared between two RVs and is defined as (Shannon, 1948; Cover and Thomas, 2006)

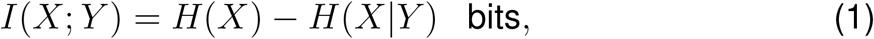

where *H*(*X*) is the entropy of *X* and *H*(*X Y*) is conditional entropy of *X* given *Y*. The mutual information can be equivalently interpreted as the amount of uncertainty remaining about *X* after observing *Y*. The mutual information, however, satisfies *I*(*X*; *Y*) = *I*(*Y*; *X*), implying that it does not describe the directional flow of information between two processes.

Unlike mutual information, the transfer entropy captures the directional flow of information between *X* (e.g., neuronal activity in the MFC) and *Y* (e.g., neuronal activity in the DS) (Schreiber, 2000; Paluš et al., 2001; Gourévitch and Eggermont, 2007; Bossomaier et al., 2016), and is defined as

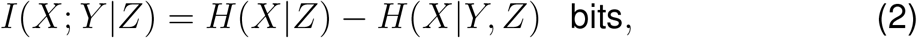

with *Z* being the history of *Y* (Figure 3). Moreover, *H*(*X | Z*) is the conditional entropy of *X* given *Z*, and *H*(*X | Y, Z*) is the conditional entropy of *X* given both *Y* and *Z*.

In this study, we test whether past neuronal activities, *Z*, give the DS the ability to extract more information from MFC signals than otherwise without them. For this purpose, we need to measure the interaction influence of *Z*. We can quantify this influence by computing the interaction information, *S*, which is defined as the extra transfer entropy remaining after subtracting the mutual information (Yeung, 1991; Gat and Tishby, 1999; Brenner et al., 2000):

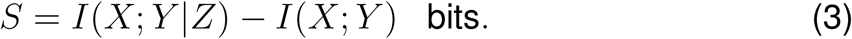

If *S >* 0, then *Z* positively influences communication rates and accordingly yields synergistic information (equation 5). Otherwise, if *S <* 0, then *Z* gives redundant information. With some algebraic manipulation, we can express equation 3 as

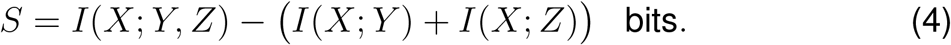

When *S >* 0, we have

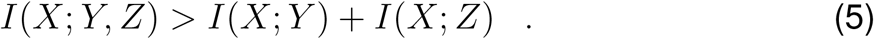

In words, equation 5 says that the whole, *I*(*X*; *Y, Z*), is greater than the sum of the parts, *I*(*X*; *Y*) + *I*(*X*; *Z*), or equivalently a synergy is present—i.e., when *Y* and *Z* work together to decode *X*, more information about *X* is obtained than if they were to work individually. As such, the interaction information reveals whether knowing *Z* enables us to extract additional information from *Y* about *X*.

Additionally, relating synergistic information to prediction can be best explained as follows: an MFC signal *X* takes ~5 ms to arrive at the DS, and past DS signals, *Z*, help decode upcoming *X* signals (i.e., *I*(*X*; *Y* |*Z*) *> I*(*X*; *Y*)).

Positive interaction information can convey an important message about the neuronal communication between the MFC and DS. When *Z* represents the history of *Y* in the immediate past, *S* is likely to be positive if the neuronal activities of the DS (*Y* and *Z*) can to some degree predict the neuronal activity of the MFC (*X*). This hypothesis is worthwhile to examine: if the hypothesis is true, it could provide us with some potential clues about how the MFC and DS share information with each other.

### Presence of Synergistic Information

Figure 4 shows rates of synergistic information (bits/sec) obtained from the neuronal activity data of 12 rats in four different trial conditions (reward versus non-reward, and left choice versus right choice). For statistical significance evaluation, we shuffled spike trains of neurons (see Materials and Methods). Here, we report rates of synergistic information 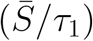 as opposed to synergistic information alone 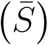 so as to put quantities in a time context. For the reason mentioned later in this section, we examined synergistic information rates for all possible combinations of reward locations (left or right spout) and outcomes (reward given or withheld). The magnitudes of the rates significantly varied from rat to rat due to certain reasons such as differences in sample sizes and average firing rates of neurons. In the majority of cases (at the p= 0.05 level), a synergistic information rate is present, that is, 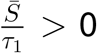. However, in Figure 4C for rodent ID #1012 and at *τ*_2_ = 10 ms a faint synergy was observed but not significant at the p = 0.05 level and therefore was discarded (and similarly, in Figure 4D, for rat #982 and at *τ*_2_ = 10 and 20 ms; as well as in Figure 4D for rat #1012 and at *τ*_2_ = 20 ms). Additionally, Figure 4 reveals that, by and large, there is a gradual increase in rates of synergistic information as more history, *τ*_2_, is included.

**Figure 4.**
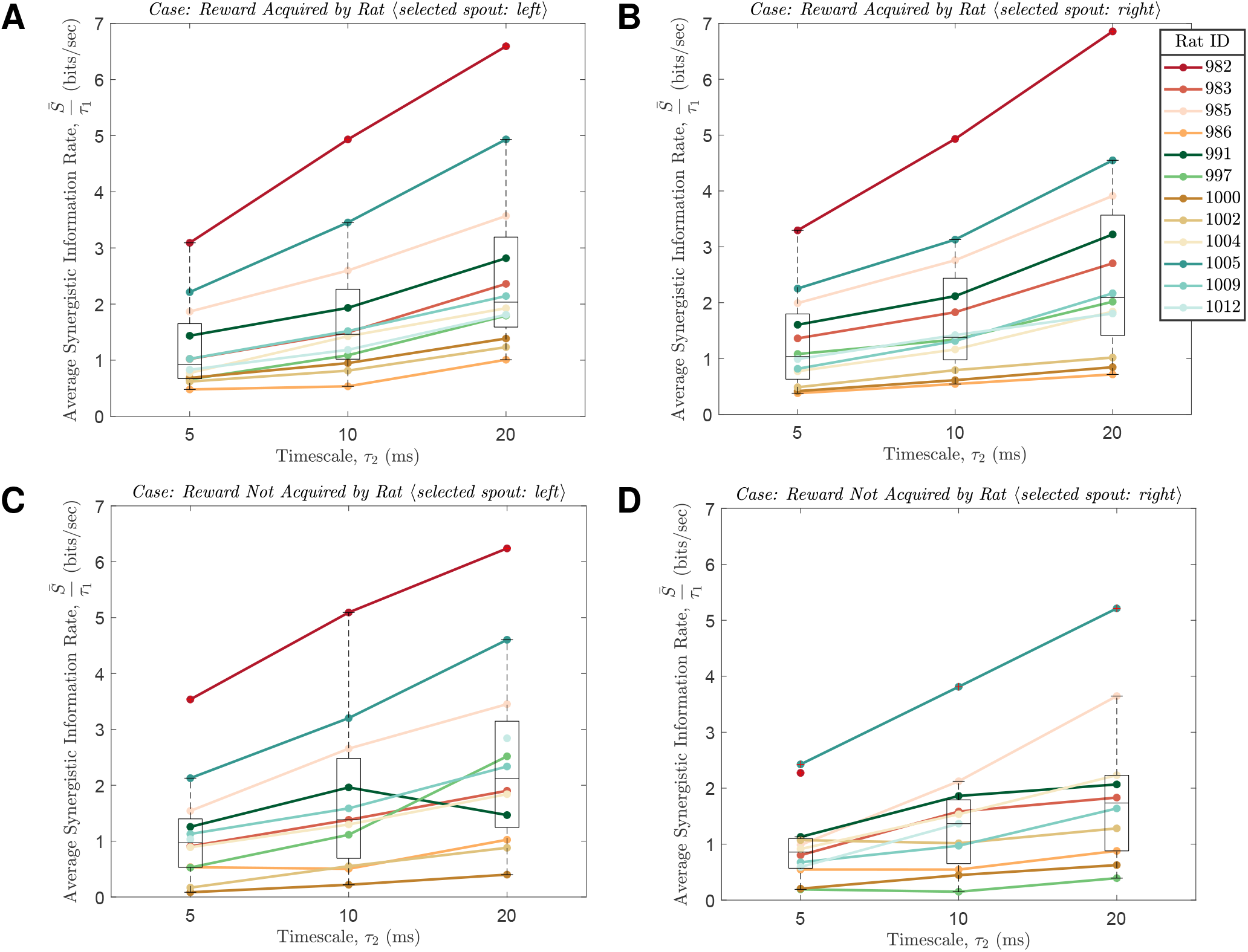
Synergistic information rates. (A) Reward acquired by rat, and the selected spout is left. (B) Reward acquired by rat, and the selected spout is right. (C) Reward not acquired by rat, and the selected spout is left. (D) Reward not acquired by rat, and the selected spout is right. In panel C, for rat #1012 at *τ*_2_ = 10 ms a faint synergy was observed but not significant at the p = 0.05 level and therefore was discarded (and similarly, in panel D for rat #982 at *τ*_2_ = 10 and 20 ms; as well as in Figure 4D for rat #1012 at *τ*_2_ = 20 ms). Here we set *τ*_1_ to 5 ms as this value is within the range of time it takes an MFC signal to reach the DS (Wilson, 1986). Additionally, we vary the value of *τ*_2_ from 5 to 20 ms to observe its influence on information rates. Boxes within plots represent the 25^th^–75^th^ percentiles and middle lines of boxes mark median values. Additionally, outlier points are beyond the ends of boxplots’ whiskers and marked with a “+” symbol.

We examined synergistic information rates during a behavioral task to rule out the possibility that the presence of synergistic information might be reward reliant or hemisphere dependent. The reason being is that in our experiment, silicon probes were inserted into rats’ left hemisphere. As such, in the behavioral task, we switched the location of rewards intermittently from left to right and vice versa, giving a total of 4 possible combinations: left/right and reward/non-reward. In our analysis, we set *τ*_1_ to 5 ms as this value is within the range of time it takes an MFC signal to reach the DS (Wilson, 1986). Additionally, we vary the value of *τ*_2_ across a range of values (5–20 ms) because a small value of *τ*_2_ may risk under-estimating the transfer entropy, while a large value of *τ*_2_ may, on the other hand, risk over-estimating the transfer entropy as a result of under-sampling the underlying probability distribution function (Bossomaier et al., 2016). We emphasize that in nearly all cases, synergistic information is present.

Finally, we test whether synergistic information in the DS is related to the sequentiality of population activities observed in DS neurons (Zhou et al., 2020). Neuronal populations in the MFC and DS often exhibit similar sequential firing patterns during behavioral tasks. Recently, a notable difference between MFC and DS activity patterns was observed in rodents performing a two-interval timing task (Zhou et al., 2020), and an outcome-based alternative choice task (in which the present data were recorded) (Handa et al., 2021). In both studies, neuronal dynamics of the DS showed a higher degree of sequentiality than those of the MFC. It is hypothesised that the higher the sequentiality of a neuronal signal, the easier it is for downstream brain areas to monitor and read out time from such signal. In the DS, monitoring time could potentially imply that it is tracking cortical inputs. However, no significant correlation was found in the present study between synergistic information rates and sequentiality across rats (Figure 5).

**Figure 5.**
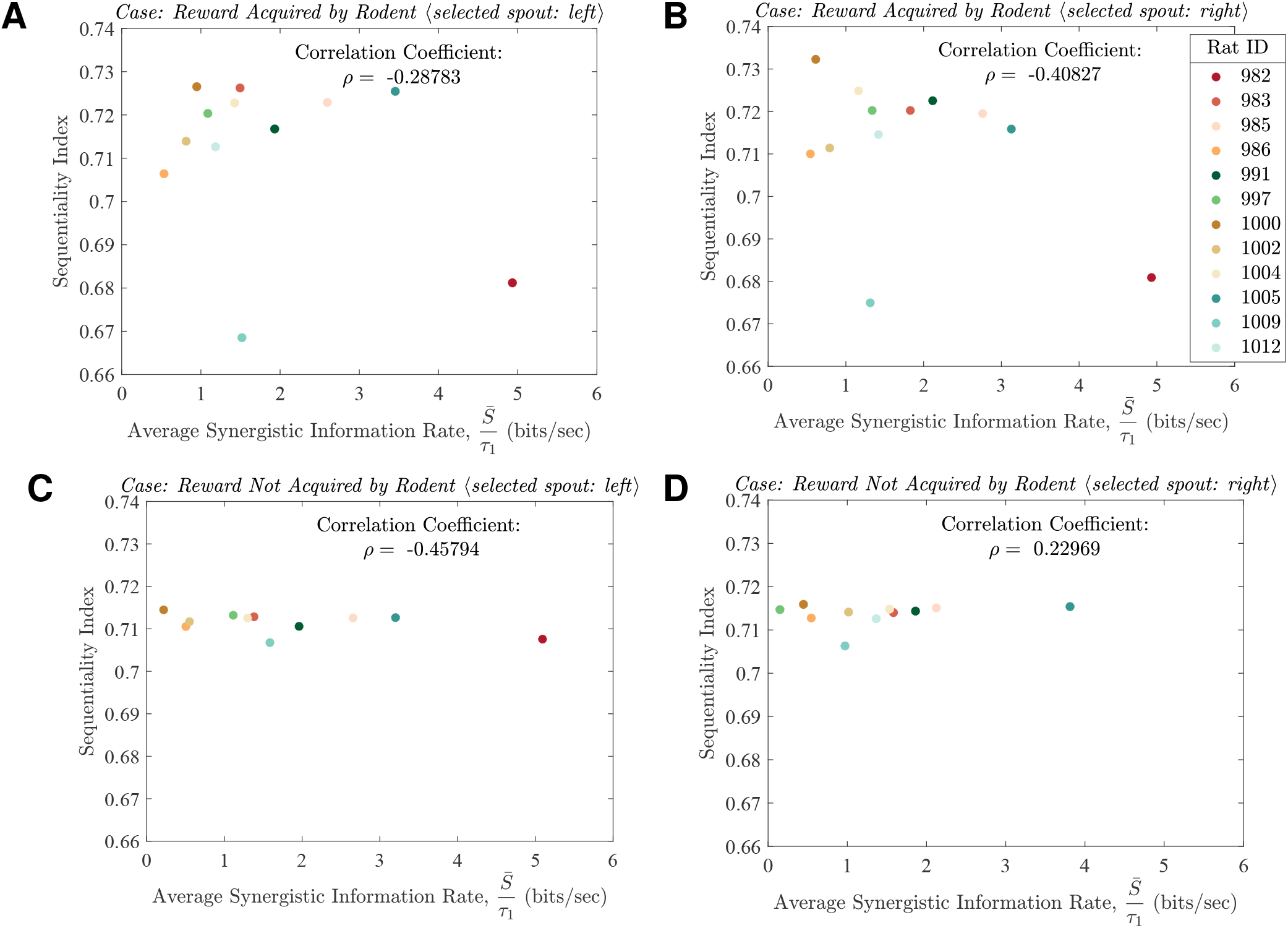
Synergistic information rates versus sequentiality index at the dorsal striatum (DS) (A) Reward acquired by rat, and the selected spout is left. (B) Reward acquired by rat, and the selected spout is right. (C) Reward not acquired by rat, and the selected spout is left. (D) Reward not acquired by rat, and the selected spout is right. In panel C, for rat #1012 a faint synergy was observed but not significant at the p = 0.05 level and therefore was discarded (and similarly, in panel D for rat #982). Here we set *τ*_1_ to 5 ms as this value is within the range of time it takes a medial frontal cortex (MFC) signal to reach the DS. Measurements are at timescale of *τ*_2_ = 10 ms, and the time-bin duration for the sequentiality index is 50 ms.

### Delinking Autocorrelation from Synergistic Information

A frequent misconception is that the inherent autocorrelation present in DS spiking activities gives rise to synergistic information and explains its predictive ability—that is to say, synergistic information is a consequence of the correlation between past and future DS spiking activities. Here we provide an example to show that this is not the case. In the example below (Figure 6), two spiking activities of the DS with the same autocorrelation give distinctly different values of synergistic information.

**Figure 6.**
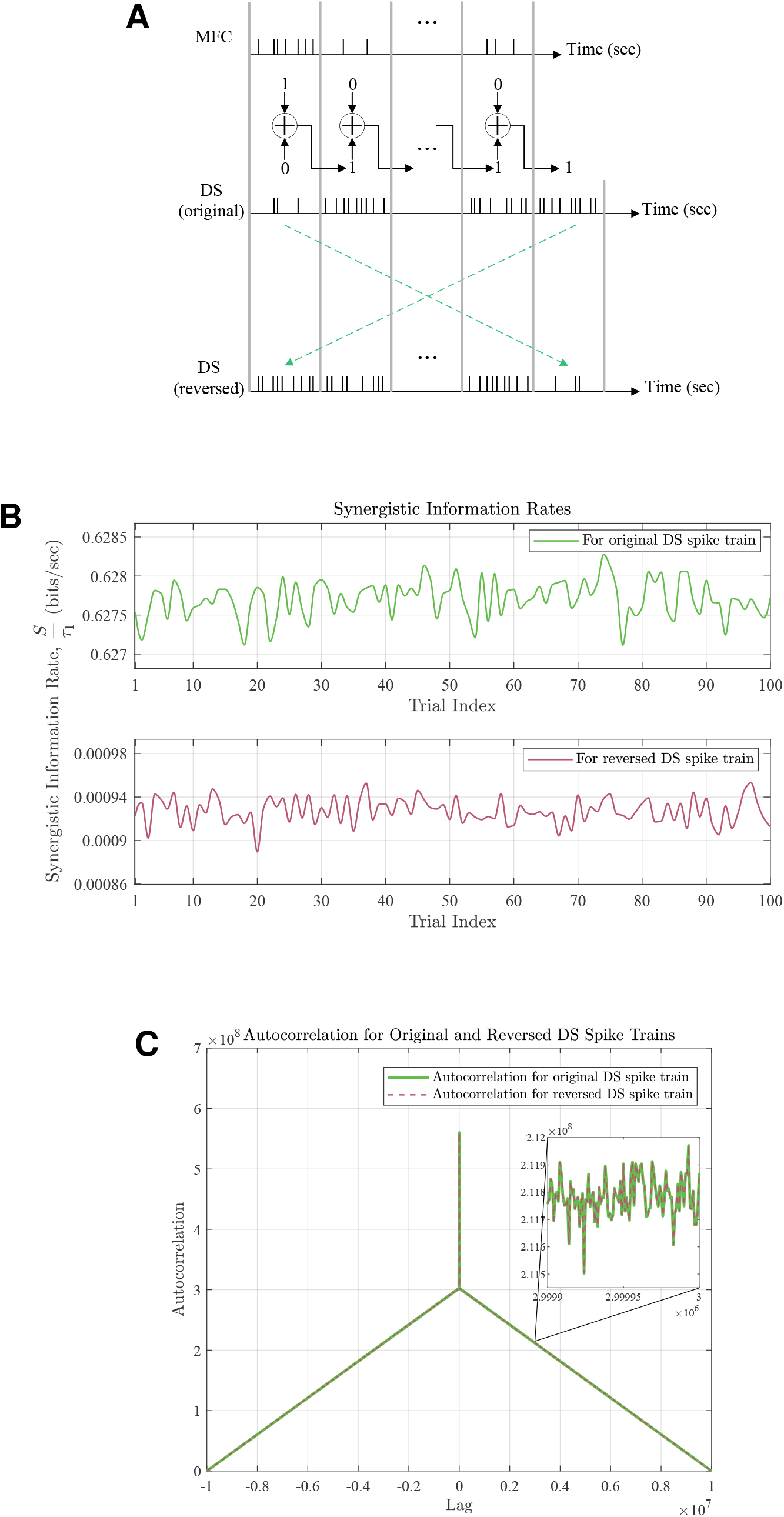
An example to illustrate the disjunction between autocorrelation and synergistic information. (A) Spiking activities of the MFC and DS. The bitstream of the MFC and DS are exclusive-ored, denoted here by the symbol ⊕. If a bit value of a timeslot is 1, then the number of neuronal spikes within the timeslot is generated according to a Poisson distribution with a mean value of *λ*_1_ = 10 (spikes/sec); otherwise, this value is set to *λ*_0_ = 0.1 (spikes/sec). (B) Synergistic information rates for the original and reversed DS spike trains. (C) Autocorrelation curve for the original and reversed DS spike trains at trial index #50.

In the illustration shown in Figure 6A, for each trial, a random bitstream of length 10^7^ is generated for the MFC with 0 and 1 being equally probable, whereas the DS bitstream is generated as follows: a future DS bit value is formed by exclusive-oring (XOR) a past MFC and DS bit with each other. If a bit value of a timeslot is 1, then the number of neuronal spikes within the timeslot is generated according to a Poisson distribution with a mean value of *λ*_1_ = 10 (spikes/sec); otherwise, this value is set to *λ*_0_ = 0.1 (spikes/sec). Additionally, in this example the duration of all timeslots are equal *τ*_1_ = *τ*_2_ = 1 (sec).

Here we consider two cases: 1) the original DS spike train is left intact 2) the original DS spike train is time-reversed. For these two cases, the MFC spike train is exactly the same. In both cases, the autocorrelation of the DS spike trains is identical. However, they give different values of synergistic information. In the first case, the amount of synergistic information is *S ≈* 0.6 bits, whereas in the second case *S ≈* 0 bits. Although there is an inherent autocorrelation in DS spiking activities, the existence of synergistic information is not necessarily a consequence of it.

## DISCUSSION

A synergistic information rate is present as past neuronal activities of the DS (*Z*) enables it to extract more information (*I*(*X*;*Y*|*Z*)) from MFC signals than otherwise without them *I*((*X*;*Y*)). What is interesting about the data is that the DS can decode information contained in MFC signals in a synergistic (*I*(*X*; *Y, Z*)) rather than an additive manner *I*(*X*; *Y*) + *I*(*X*; *Z*). Our results suggest that past DS activities have a useful role in the MFC-DS communication network because they enhance the communication rate between the two brain regions. Whether past striatal activities contribute to the cortico-striatal information transfer has not been previously explored.

Searching for synergistic information in a neural code has been undertaken and found in a variety of settings. In blowflies, synergistic information was detected in H1-neurons. It was observed that two action potential spikes together, across time, carry more than twice the amount of information of individual spikes (Brenner et al., 2000). In salamanders and guinea pigs, a similar synergy was detected in a combinatorial code of neurons, whereby spiking and silent neurons jointly produce synergistic information in a neural pattern (Schneidman et al., 2011). Likewise, synergistic information in patterns of neuronal activities at the population level was further observed in macaque monkeys (Osborne et al., 2008). The previously described studies examined synergistic information in neuronal representations within a single brain region. What sets our work apart from prior research work is that we examine the synergy forming across brain regions—the MFC and DS, the critical brain regions implicated in decision-making. Our results indicate that synergistic information is also present in neural codes for cross-area communication.

The interaction between the MFC and DS has been gaining attention. Results from earlier studies revealed that ramping activities of the MFC and DS are likely to represent temporal signals that encode the passage of time (Emmons et al., 2017). Additionally, it was demonstrated that optogenetically stimulating MFC-to-DS axonal projections increase time-dependent ramping in the DS (Emmons et al., 2019). In these studies, the linear relationship between the MFC and DS was studied. Our results are complementary to these studies in that we examined a nonlinear relationship between the MFC and DS by means of information-theoretic tools. As such, a novel interaction is observed, namely, the emergence of synergistic information once past neuronal activities of the DS are considered.

We propose that synergistic information, at least partially, emerges from the computational effort of the DS to predict neuronal activity in the MFC in the immediate future—within the milliseconds range. We consider that this predictive role of the DS is unrelated to reward prediction. A body of evidence shows that the striatum represents error signals of reward predictions (Oyama et al., 2010; Stalnaker et al., 2012) that modify the efficacy of neuronal signal transmission in the cortico-striatal pathway (Cools, 2011; Morita et al., 2012). The timescale of this prediction is behaviorally relevant in the range of seconds, which is much longer than the millisecond range timescale of synergistic information at the DS. Additionally, no particular differences between rewarded and unrewarded trials in terms of rates of synergistic information were observed in our experiment.

Synergistic information gained by past DS activities is uncorrelated with the degree of sequentiality of the population activity in the DS, suggesting that monitoring time and predicting cortical states are not conclusively related to each other in the present study. This may simply reflect the fact that unlike the interval timing task (Zhou et al., 2020), the present outcome-based choice task does not require an accurate estimation of time. However, we also noticed that the sequentaility index used in this and previous studies is not robust against noise. The computational relationship between synergistic information and high sequentiality in the DS and the behavioral implications of these observations require further clarifications.

From an energy perspective, synergistic information improves metabolic efficiency. In recent years, there has been a particular interest in studying the energy efficiency of the mammalian brain. One line of research indicates that the mammalian brain is structured in such a way as to maximize metabolic efficiency (Harris et al., 2012, 2015, 2019; Yu and Yu, 2017)—measured as bits transmitted per energy expended to generate an information-carrying action potential. In other words, the brain is tuned to communicate economically in terms of energy expended to transmit information. Our results are in line with the metabolic efficiency viewpoint because past neuronal activities enable the DS to extract more information contained in MFC signals than without them. This, as a result, increases the metabolic efficiency of the cortico-striatal circuit: more bits extracted from a neuronal signal gives a higher metabolic efficiency. Additionally, it is useful to mention that the efficient use of metabolic energy is a widely accepted assumption behind the development of the so-called predictive coding hypothesis for hierarchical cortical computing (Rao and Ballard, 1999; Friston, 2005, 2010), although counter-arguments have also been given (Keller and Mrsic-Flogel, 2018).

A suggested hypothesis for the function of the basal ganglia proposes that the striatum selectively regulates (gates) cortical information. When the DS regulates cortical information, task-relevant information has to be extracted from MFC signals and subsequently transmitted to downstream brain regions. Such processing of cortical signals will filter out information unnecessary for controlling voluntary movements (DeLong, 1990; Ding et al., 2010; Tecuapetla et al., 2016; Klaus et al., 2019). Additionally, filtering at the DS is more likely to decrease rather than increase the amount of information extracted from the MFC signals. Our results, however, suggest that more information flows from the MFC to DS when past neuronal activities of the DS are considered than without them (i.e., *I*(*X*; *Y Z*) *> I*(*X*;|*Y*) or equivalently *S >* 0, equation 3). Although the gating hypothesis and the existence of synergistic information at the DS are not mutually exclusive, they do not harmonize with each other.

A shortcoming of our study is that we only observed a small area of the MFC and DS using two probes. We used a retrograde tracer to confirm the projection from the recording site at the MFC to its corresponding recording site at the DS. However, a large-scale experiment covering a wide area of both the MFC and DS is necessary to solidify our results. Nonetheless, all the rats examined in this study consistently supported the significant levels of synergistic information at the DS. We, therefore, believe that our results are not mere noise arising from the limited sampling of MFC and DS neuronal signals but indicate a distinct predictive function operating in the frontal cortex-striatal pathway on short timescales.

## MATERIALS AND METHODS

### Animal preparation

All procedures followed in this study were carried out in accordance with the Animal Experiment Plan, which was reviewed and approved by the Animal Experiment Committee of RIKEN. A total of 20 Long-Evans rats (6 weeks, male, 200 – 220 g, Japan SLC Inc.) were used in our experiment. In-depth details of our methods are available in our recent report (Handa et al., 2021). In the present study, we analyse the data obtained from our recent work (Handa et al., 2021).

### Behavioral task

We used a customized multiple-rat training system (Isomura et al., 2009; Handa et al., 2021) to train several rats in parallel to learn the spout-selection task and gave them saccharin-water (0.1%, 15*µ*l) as a reward. We analysed the neuronal activity of 12 rats out of the 20 we trained because they provided a satisfactory number of units from both the MFC and DS.

Figure 1A provides a schematic illustration of a single test trial. A trial starts with a 3 kHz (1 sec) tone. Then, a random delay (0.7–2.3 sec) is introduced; if a rat licks a spout during this period, the trial is aborted. The random delay is followed by a 10 kHz (0.1 sec) “Go” signal. After this signal, a rat selects either a left or right spout. Then, a second random delay (0.3–0.7 sec) is introduced before a fluid reward is either delivered or withheld; if a rat selects a correct spout location, a reward is delivered for a period of 4 sec. On the other hand, if a rat selects an incorrect spout location, a reward is withheld; the duration of this non-reward event is 5 sec. After a reward/non-reward phase is complete, a new trial begins.

Figure 1B provides an example from the behavioral experiment. First, a rat selects either a left or right spout. Then, a reward is delivered if the rat selects a correct choice. A choice is declared correct if the selected spout location matches the location of a reward block. Otherwise, the choice is declared incorrect, and a reward is withheld. The location of the rewarding spout was fixed during a trial block and was reversed after a rat accumulated a total of 10 rewards. During the experiment, we provide no sensory feedback to the rats regarding the location of the reward.

### Electrophysiological recordings from the MFC and DS

Figure 3 shows an illustration of the electrophysiological recording. We inserted the pair of silicon probes (32-channels, 4 shanks separated 0.4 mm apart) that have tetrode-like electrodes (A4*×*2-tet-7/5 mm 500-400-312, NeuroNexus Technologies) as follows:

- The first silicon probe was inserted vertically (depth from the pia mater: 1.2 mm) into the MFC (+2.4 – 4.2 mm from the Bregma and 1.0 – 1.4 mm to the midline).
- The second silicon probe was inserted at an angle of 6°posteriorly into the DS through a cranial window (+0.6 – 1.0 mm from the Bregma and 1.0 – 1.4 mm to the midline).

### Retrograde tracing

To confirm that the recording site at the MFC projects to that of the DS, we injected a retrograde tracer (Fluoro-Gold) into the DS 3 days before a first electrophysiological recording session and checked the position of labelled neurons *post hoc*. Figure 2 shows fluorescent images obtained after a recording experiment. We observed that corticostriatal projection neurons at the MFC are labelled with fluoro-gold after injecting the retrograde tracer at the DS. We collected images with a fluorescent microscope (Olympus AX70, Tokyo, Japan).

### Measurements and calculations

Neuronal signals of our experiment are analysed from the moment a rat first licks a spout (after the onset of a “Go” signal) to the end of a trial (Figure 1A). During this period, the probabilities of events and synergistic information are computed as follows:

- ***Probabilities:*** Figure 7 provides an illustration of the random variables (RVs) and time-windows considered herein. Let *X* be an RV that counts the number of spikes generated by all MFC neurons within a time-window of duration *τ*_1_. Likewise, let *Y* and *Z* be RVs that count the number of spikes generated by all DS neurons within a time-window of duration *τ*_1_ and *τ*_2_, respectively. Now let us describe how outcomes of RVs are obtained. In Figure 3, all three time-windows jointly slide by increments of *τ*_1_ to obtain an event (*x, y, z*), where *x, y*, and *z* are outcomes of *X, Y*, and *Z*, respectively. Namely, *x, y*, and *z* are the number of spikes observed in time intervals [*nτ*_1_, (*n* + 1)*τ*_1_], [(*n* + 1)*τ*_1_, (*n* + 1)*τ*_1_ + *τ*_2_], and [*nτ*_1_, (*n* + 1)*τ*_1_], respectively; where *n ∈* {0, 1, 2, …}. Probabilities are estimated by the relative frequency of occurrence of events. For instance, *p*(*x, y, z*) is the relative frequency of occurrence of the event (*X* = *x, Y* = *y, Z* = *z*).
- ***Synergistic information:*** The amount of synergistic information is computed using equation 3, where *I*(*X*; *Y*|*Z*) is the transfer entropy (Schreiber, 2000; Paluš et al., 2001; Gourévitch and Eggermont, 2007; Bossomaier et al., 2016) and *I*(*X*; *Y*) is the mutual information (Shannon, 1948; Cover and Thomas, 2006), and are calculated as follows:

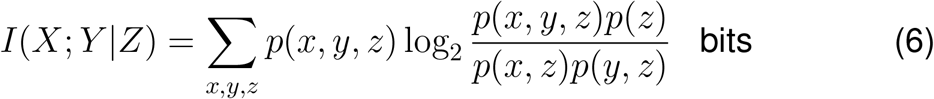

and

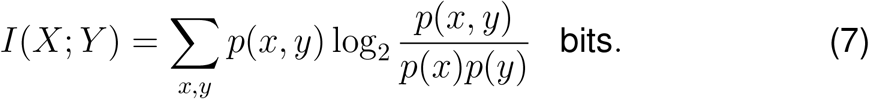

**Figure 7.**
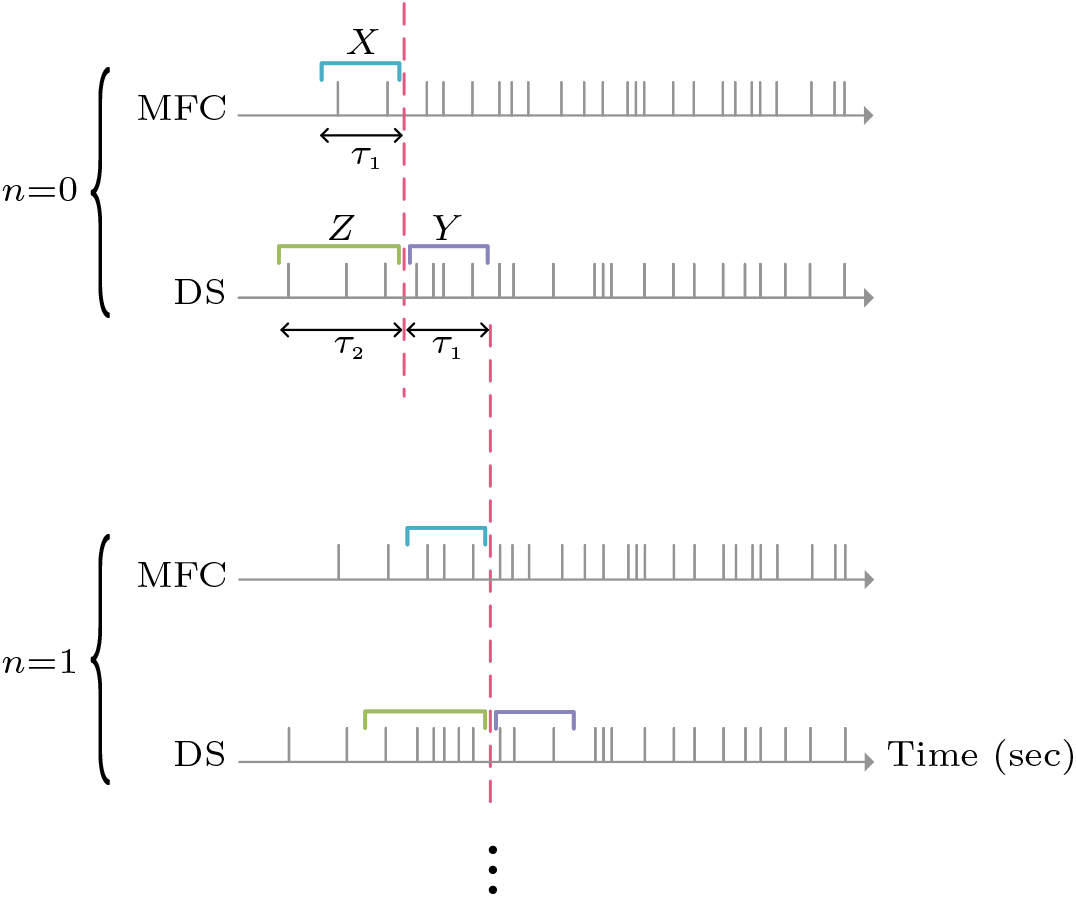
An illustration of the random variables (RVs) and time-windows used for synergistic information calculations. The medial frontal cortex (MFC) spike train displays the collective firing of all MFC neurons, rather than the firing of an individual MFC neuron—and likewise for the dorsal striatum (DS) spike train. RV *X* counts the number of spikes generated by all MFC neurons within a time-window of duration *τ*_1_. Similarly, RVs *Y* and *Z* count the number of spikes generated by all DS neurons within a time-window of duration *τ*_1_ and *τ*_2_, respectively. In this particular example, at instance *n* = 0, the outcomes (i.e., the number of spikes) of the RVs are 2, 4, and 3, for *x, y*, and *z*, respectively. At *n* = 1 all three time windows shift to the right by *τ*_1_, and the outcomes of the RVs are 3, 3 and 6, for *x, y* and *z*, respectively.

### Statistical significance

This section is based in part on (Wibral et al., 2014; Bossomaier et al., 2016; Timme and Lapish, 2018; Timme et al., 2020). We test the statistical significance of our results by the following procedure:

- First, for each experimental trial, we generate null (surrogate) data of the MFC by the following method: consider the illustration shown in Figure 8. The MFC spike train of each experimental trial is divided into two parts by a random number, *t*^*∗*^, drawn from a uniform distribution *U* (*t*_start_, *t*_end_), where *t*_start_ and *t*_end_ are the start an1d end time of the MFC spike train, respectively. Next, the two MFC parts are swapped to create null data. Ideally, this makes the MFC and DS spike trains independent of each other—effectively disconnecting the MFC from the DS in terms of shared information while preserving much of the statistical properties of the original MFC spike train. This randomization procedure is repeated *N* = 10^4^ to form null data for each experimental trial. Additionally, the DS spike train was left unrandomized to preserve the statistical relationship between *Y* and *Z* (Bossomaier et al., 2016).
- Second, using the MFC null data generated in the previous step along with the intact DS spike train, we use equations 6 and 7 to compute the set *I*_s_(*X*; *Y*) and *I*_s_(*X*; *Y* |*Z*) (the null hypothesis versions of *I*(*X*; *Y*) and *I*(*X*; *Y* |*Z*), respectively).
- Third, the p-value of *I*(*X*; *Y*) is estimated as the fraction of *I*_s_(*X*; *Y*) values that are greater or equal to *I*(*X*; *Y*). Likewise, the p-value of *I*(*X*; *Y* |*Z*) is estimated as the fraction of *I*_s_(*X*; *Y* |*Z*) values that are greater or equal to *I*(*X*; *Y* |*Z*).

**Figure 8.**
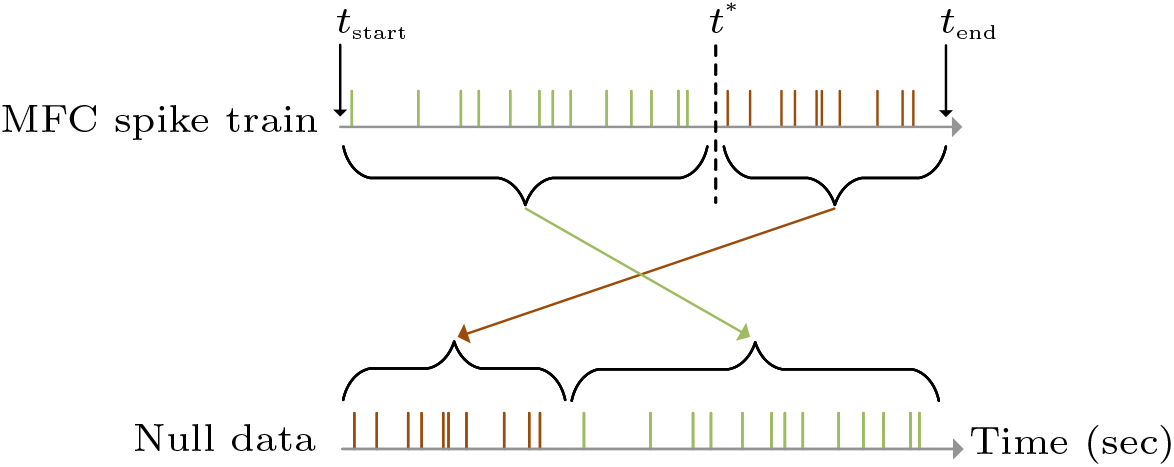
An illustration to describe how null data is generated. Here, *t*^*∗*^ is a random number drawn from a uniform distribution *U* (*t*_start_, *t*_end_), where *t*_start_ and *t*_end_ are the start and end time of the medial frontal cortex (MFC) spike train, respectively. The value of *t*^*∗*^ divides the MFC spike train into two parts, which are swapped to form null data.

### Calculation remarks

- After we calculate *I*(*X*; *Y*) and *I*(*X*; *Y*|*Z*), and their associated p-value for each experimental trial, we take the average of all *S* values for which both *I*(*X*; *Y*) and *I*(*X*; *Y*| *Z*) have a p-value ≤ 0.05. We denote this average by 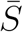.
- Consider Figure 1B. If a trial before an incorrect choice is as well incorrect, then it is discarded in our calculations. For example, trial #196 in Figure 1B was discarded while trial #197 was retained. Such events are not uncommon in our experiment, and we discard them to preserve consistency.

## ACKNOWLEDGEMENTS

The authors would like to gratefully thank Thomas Burns, Milena M. Carvalho, Ruxandra Cojocaru, and Yukiko Goda for their assistance and discussions. This research was made possible by the Japan Society for the Promotion of Science (JSPS) research grant (Grant Number: 20K07716 to TH), Hiroshima University Start-up Grant, and RIKEN grant. Illustrations herein were primarily created with BioRender.com

## AUTHOR CONTRIBUTIONS

T.F. and I.A. designed the project. T.H. performed both the behavioral and electrophysiological experiments, and I.A. analysed the neuronal data. All authors participated in writing the manuscript.

